# Thalamic blood flow and EEG frequency band power during hyperventilation in idiopathic generalized epilepsy

**DOI:** 10.1101/2024.03.26.586742

**Authors:** Justine Debatisse, Siti N Yaakub, Enrico De Vita, Geoff Charles-Edwards, Sami Jeljeli, James Stirling, Anthime Flaus, Joel Dunn, Michalis Koutroumanidis, Alexander Hammers, Colm J McGinnity

## Abstract

Absence seizures in idiopathic generalised epilepsy (IGE) involve thalamo-cortical circuits. Hyperventilation (HV) is a standard technique to trigger epileptiform discharges and absence seizures in IGE. HV also increases electroencephalography (EEG) delta band power and decreases global cerebral blood flow (CBF). The relationships between HV, EEG band power, and regional CBF have not been investigated in the same patients at the same time.

We compared the effects of hyperventilation between 13 individuals with IGE and 18 healthy controls on thalamic CBF assessed with pseudo-continuous Arterial Spin Labelling (pCASL) simultaneously to EEG frequency band power during three periods of hyperventilation interleaved with three periods of rest, normalised to each participant’s pre-HV values.

Pre-HV, there were no differences between patients and controls. During the three rest periods combined, patients had higher normalised respiratory rates but no difference in CBF or EEG band power compared with controls.

During HV, CBF decreases were similar in controls and IGE patients in cortical gray matter (37.1±1.3% in controls, 37.9±2.0% in patients) and basal ganglia (35.6±2.0% in controls, 33.3±2.3% in patients). In the thalamus, CBF decreased by 35.5±1.9% in controls, but only by 27.1±2.5% in patients (p=0.011). Delta band power increased by 42% in patients, but only by 27% in patients (p<0.045).

Both controls’ and patients’ thalamic CBF significantly decreased as a function of respiratory rate, but the relationship was weaker in patients (rho –0.38 [0.95 confidence interval lower limit −0.629; upper limit −0.059] vs –0.51 [-0.698; −0.255] in controls). Delta power correlated with thalamic CBF only in controls (rho −0.21 [-0.388; −0.017]) but not in patients (rho −0.1 [-0.345; 0.157]). EEG power and respiratory rate were not significantly correlated in either group.

This first study of the interrelationships between HV, blood flow, and EEG band power suggests that regional thalamic abnormalities of blood flow regulation, or a neurophysiological abnormality leading to smaller reactive blood flow changes, may underlie HV-associated EEG activation.

**Graphical abstract text:** In this first investigation of combined hyperventilation, blood flow and EEG band power measurements in idiopathic generalised epileptic patients and healthy subjects, we revealed that localized thalamic blood flow regulatory problems or neurophysiological disorders caused minor reactive blood flow variations and may cause hyperventilation-associated EEG activation.

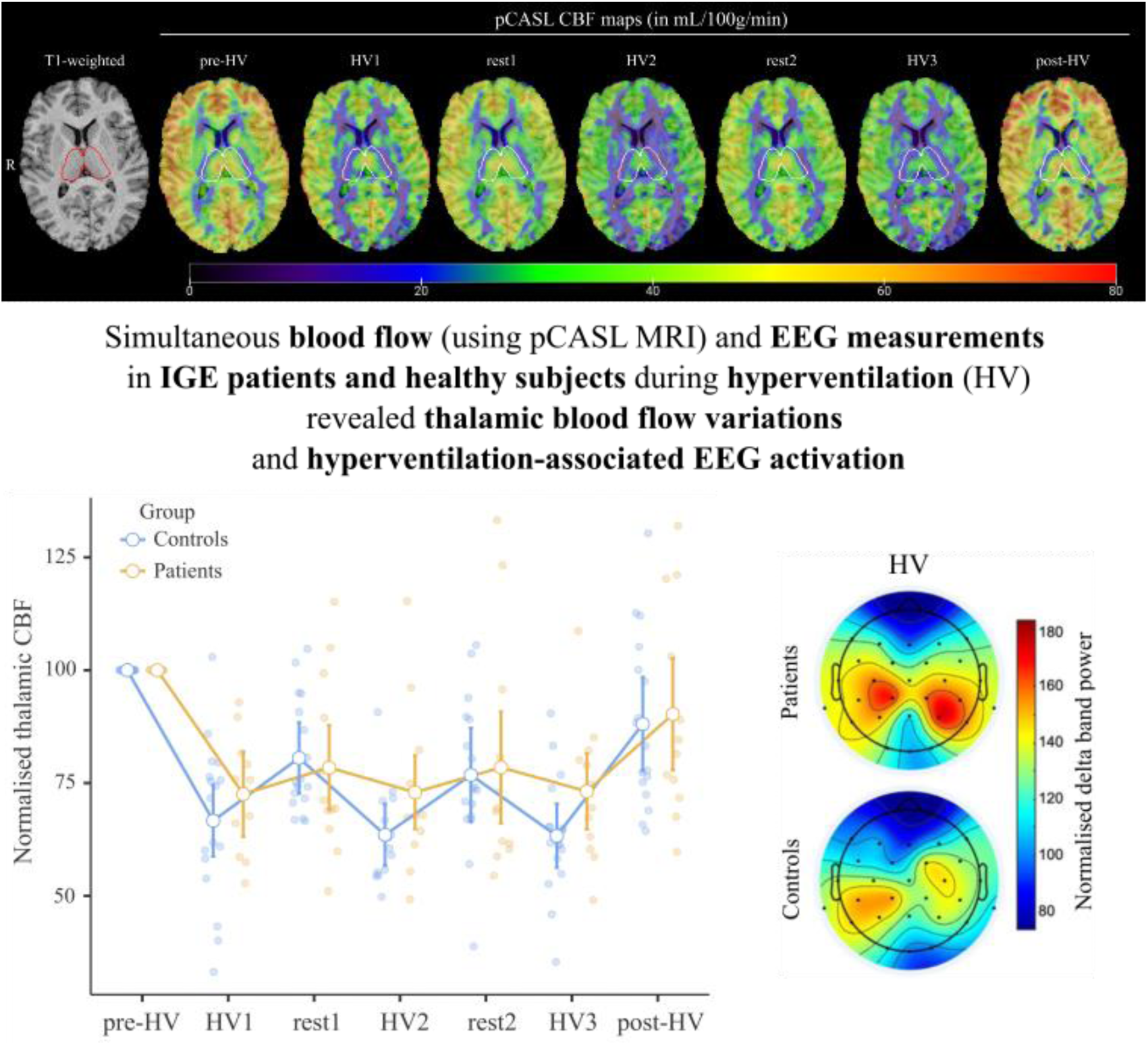

## Introduction

The International League Against Epilepsy has proposed the term “idiopathic generalized epilepsies” (IGEs) is reserved for a distinct subgroup of genetic generalized epilepsies that includes childhood and juvenile absence epilepsies (Hirsch, French, Scheffer, Bogacz, Alsaadi, et al., 2022). Electroencephalographic findings in the IGEs are typified by generalized spike-wave and/or polyspike-wave electroencephalographic discharges that may activate on provocation (Hirsch, French, Scheffer, Bogacz, Zhou., et al., 2022).

Hyperventilation (HV), i.e. voluntary over-breathing unrelated to exercise, is a provocation technique that is widely used clinically to trigger epileptiform discharges and absence seizures in individuals with suspected IGE (Foerster, 1924; Rosett, 1924). HV induces a decrease in the partial pressure of carbon dioxide (pCO_2_) in the blood i.e. hypocapnia that persists for several minutes after the cessation (Blinn & Noell, 1949; Kety & Schmidt, 1946). Hypocapnia promotes the association of H^+^ and bicarbonate ions in the blood, resulting in an elevation in pH (potential of hydrogen) – respiratory alkalosis.

Blood vessels constrict in response to an elevation in pH in order to maintain homeostasis (Elliott & Jasper, 1949). Accordingly, hyperventilation has been associated with marked reduction in cerebral blood flow (CBF; e.g. nearly one-third in (Kety & Schmidt, 1946); up to 27% in (Pavilla et al., 2018), up to 37% in (Ito et al., 2002).

The normal electroencephalographic (EEG) response to hyperventilation is characterised by an increase in overall amplitude (i.e. power) and decrease in mean frequency (Berger, 1934; Gibbs et al., 1968). For example, Van der Worp et al. reported increases in total power of up to 398 μV^2^ in association with hyperventilation and decreases in the mean frequency of up to 3.9 Hz; blood flow velocity in the middle cerebral artery was also reduced by up to 53±12% (Van der Worp et al., 1991). HV produced a more profound and sustained decrease in pCO_2_ in the jugular vein in individuals with absence epilepsy than in healthy controls (Nims et al., 1940). Twenty children with epilepsy, including four with absence epilepsy, had lower mean EEG frequencies than controls during HV (Yamatani et al., 1995). They also had a greater decrease in global CBF assessed via Doppler ultrasound of the right common carotid artery in the first minute of hyperventilation.

The thalamus is a region of interest implicated in the pathophysiology of absence seizures in IGE Simultaneous EEG and functional MRI studies have found increased activity in thalamocortical circuits during unprovoked spike and wave discharges, suggesting the involvement of the thalamus in the generation or spread of discharges in IGE (Aghakhani et al., 2004; Berman et al., 2010; Gotman et al., 2005). Studies looking at the dynamic evolution of unprovoked generalised spike and wave discharges found that they are often driven by a change in activity in the cortex followed by an increase in activity in a network of thalamocortical regions (Moeller et al., 2008; Tangwiriyasakul et al., 2018; Vaudano et al., 2009). A study investigating CBF distributions using arterial spin labelling (ASL) found deficient interictal cerebral perfusion in the thalamus, upper midbrain, and left cerebellum (Sone et al., 2017), suggesting that thalamic abnormalities persist outside of seizure periods. Involvement of the thalamus is also suggested by diffusion MRI, magnetic resonance spectroscopy (Fojtiková et al., 2006; Kabay et al., 2010), [^15^O]H_2_O positron emission tomography (Prevett et al., 1995), and voxel-based morphometry imaging studies (Pardoe et al., 2008), as well as extensive animal model literature (for a review, see (Depaulis & Charpier, 2018)).

The aim of this work was to study the interplay between repeated hyperventilation, thalamic cerebral blood flow (CBF), and electroencephalography (EEG) frequency band power between individuals with IGE and healthy controls with simultaneous EEG-MRI.

## Materials and methods

### Participants

Participants were recruited as part of a PET-MR-EEG study on dopamine neurotransmission. The primary inclusion criteria for the epilepsy group were: 1) IGE with very frequent and/or hyperventilation-inducible absence seizures; and 2) age 16 years or more.

Exclusion criteria were 1) history of neurological or psychiatric disease that was likely to influence dopaminergic neurotransmission (e.g. Parkinson’s, schizophrenia); 2) regular or recent medication intake that was likely to influence dopaminergic neurotransmission, 3) any contraindication for undergoing MRI; 4) regular or recent use of a recreational drug; 5) claustrophobia; 6) participation in a trial involving medicinal products within the preceding three months; 7) participation in research involving ionising radiation within the preceding year; 8) inability to provide informed consent; 9) history of chronic or recent cardiac, cerebrovascular, moyamoya or pulmonary disease, or of sickle cell disease/trait. Antiseizure medication was allowed for the patients.

Thirteen patients (median ± interquartile range age 32.0 ± 18.0 years; four males) were included; they were recruited from outpatient epilepsy clinics at Guy’s and St Thomas’ and King’s College Hospital NHS Foundation Trusts (both London, UK). The diagnosis of IGE-absences was based on history and seizure semiology, as well as interictal and ictal electroencephalography (EEG) recordings and brain magnetic resonance (MR) imaging. The electroclinical features that we deemed suggestive of absence seizures included brief episodes of behavioural arrest and impairment of awareness, accompanied by generalised spike-and-wave or polyspike-and-wave discharges at approximately 3 Hz. All individuals with IGE-absences had normal conventional brain MR.

Eighteen healthy controls (30.0 ± 14.3 years; five males) without history of epilepsy were also recruited via local advertising. They had no regular or recent medication intake; exclusion criteria were otherwise the same as for patients.

### Urinary drug screening

Participants underwent a urine drug cassette test (ALLTEST 12 panel (7+5) Workplace NPS; Hangzhou AllTest Biotech Co., Ltd; Hangzhou, China; http://www.alltests.com.cn/EN/) to check for the following substances approximately one hour before the PET scan: amphetamine; benzodiazepines; cannabis; cocaine and crack cocaine; heroin and all opiates; K2/synthetic cannabinoids; ketamine; MDMA/ecstasy, MDPV/bath salts; MEP/mephedrone; methadone; methamphetamine. Two patients who were prescribed as-needed benzodiazepines and had taken a dose 2 and 7 days previously, respectively, tested positive for benzodiazepines. One patient (benzodiazepines) and one control (morphine) had unexplainedly positive results.

### Ethical considerations

The West Midlands – Coventry & Warwickshire Research Ethics Committee granted ethical approval for this study (16/WM/0364). All participants provided written informed consent in accordance with the Declaration of Helsinki.

### MR and EEG data acquisition overview

The scans were acquired at the King’s College London and Guy’s and St Thomas’ PET Centre on a Biograph mMR PET-MR scanner ((Delso et al., 2011); Siemens, Erlangen, Germany). Each participant had a 60-minute EEG-MR scan that included a 3D T1-weighted acquisition and a pseudo-continuous arterial spin labelling (pCASL) sequences (Dai et al., 2008; Wu et al., 2007), see below.

Thirty-two channel EEG was acquired throughout the scan, during which the participants also wore a belt to monitor their respiratory rate. EEG data was acquired using an MR-conditional EEG system capable of acquiring data within the MR environment. Participants were fitted with a BrainCap MR EEG cap (Brain Products GmbH, Gilching, Germany) with 32 Ag/AgCl electrodes arranged according to the extended international 10-20 system with the reference electrode placed between Fz and Cz and the ground between Fz and Fpz. The electrocardiogram (ECG) was recorded at a sampling frequency of 5 kHz using the BrainVision Recorder software (Brain Products).

The individuals with epilepsy were closely observed for evidence of seizures throughout the scan (none detected).

### Hyperventilation task

At approximately 24 minutes into the scan, participants performed auditory-cued hyperventilation (HV) at a rate of ∼24 breaths per minute, as per clinical EEG practice. The first 18 participants (6 with epilepsy, 12 controls) performed three blocks of three-minute HV, each followed by two minutes of free breathing. The hyperventilation protocol was revised for the final 13 participants (final seven participants with epilepsy and six controls) in an attempt to elicit generalized spike-and-slow-wave discharges more reliably in participants with epilepsy. These 13 participants performed two blocks of four-minute HV, each followed by three minutes of free breathing, followed by a final block of two-minute HV.

Pseudo-continuous arterial spin labelling (pCASL) data were acquired from four minutes before until approximately eight minutes after the task. Figure 1 illustrates the task protocol and data acquisition.

**Figure 1.**
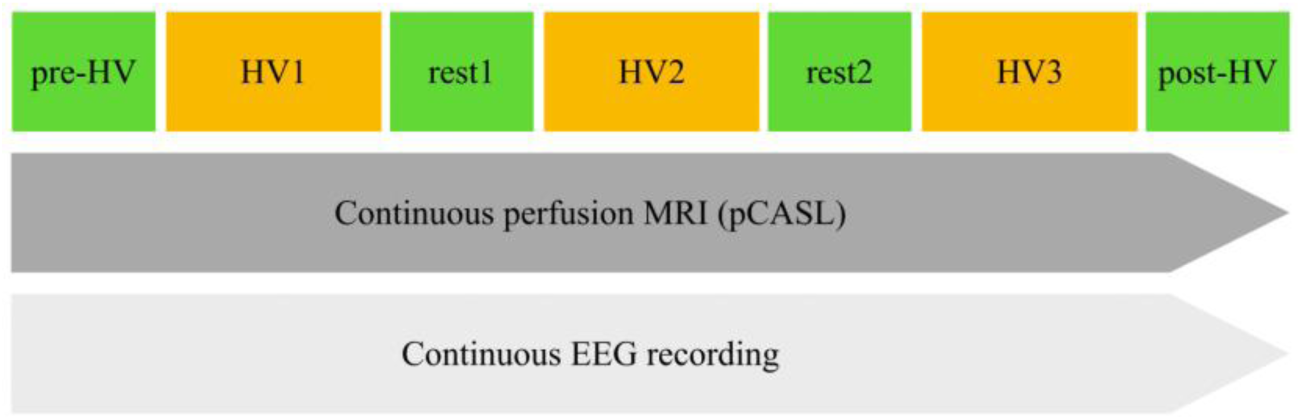
Description of the simultaneous EEG recording and perfusion MRI (pCASL) imaging during the hyperventilation-rest task, starting ∼24 minutes into the 60-minute scan. HV blocks lasted 4 minutes for n=13 participants and 3 minutes for n=18 participants. Rest blocks lasted 3 minutes for n=13 participants and 2 minutes for n=18 participants. EEG: electroencephalography: HV: hyperventilation; MRI: magnetic resonance imaging; pCASL: pseudo-continuous Arterial Spin Labelling

### Structural MR data acquisition and processing

The T1-weighted structural data was acquired with a 3D MPRAGE sequence with: repetition time (TR)/echo time (TE) = 1700/2.19 ms, inversion time (TI) = 900 ms, flip angle 9 degrees, acquired voxel size = 1.0 × 1.0 × 1.0 mm. The brain was exracted with pincram ((Heckemann et al., 2015); https://github.com/soundray/pincram) and segmented into 95 anatomical regions from the Hammers Brain Atlas Database ((Faillenot et al., 2017; Gousias et al., 2008; Hammers et al., 2003; Wild et al., 2017); https://www.brain-development.org/atlases) using MAPER ((Heckemann et al., 2010); https://github.com/soundray/maper). Right and left thalamus were merged into one single label. The basal ganglia label included left and right caudate nucleus, pallidum, nucleus accumbens, putamen and substantia nigra. Tissue-class segmentation of the T1 images was performed via Unified Segmentation as implemented in SPM12 (https://www.fil.ion.ucl.ac.uk/spm/; The Wellcome Centre for Human Neuroimaging, UCL Queen Square Institute of Neurology, London, UK). The output grey matter image was thresholded at 0.5 probability, and then multiplied with the MAPER output atlas to produce an individualised grey matter masked regional cortical atlas where all supratentorial labels (74 labels out of 95, without ventricles, corpus callosum, brainstem, cerebellum, basal ganglia and thalamus) were merged into one single label called cortical gray matter.

### EEG data processing

The EEG data were processed using BrainVision Analyzer (v2.1, Brain Products GmbH, Gilching, Germany). Data recorded whilst the MR sequences were not running were downsampled to 500Hz, filtered (high-pass 1 Hz, low-pass 70Hz, notch 50Hz), and corrected for cardioballistic artifacts after filtering the ECG electrode data (1 Hz, 7.5Hz). Data recorded during pCASL sequences were additionally corrected for MR gradient artifacts.

The processed EEG data acquired from participants with epilepsy were carefully reviewed for evidence of epileptiform discharges in standard and bipolar montages (by CJM and MK).

Data were then segmented into hyperventilation and rest blocks according to the hyperventilation protocol for each participant, before being further processed and analysed using custom scripts (Yaakub et al., 2020) with tools from the FieldTrip toolbox (Oostenveld et al., 2011) implemented in MATLAB (R2020b, The MathWorks Inc., Natick, MA). The first 10 s and last 10 s of EEG data in each block were removed and the remaining data split into trials of 10 s length. Data were re-referenced to the average of all channels except for Fp1, Fp2 (to exclude eye movement artifacts from the average) and ECG, the linear trend was removed from each trial and baseline correction performed. Artifacts in the form of large spikes, muscle activity (in channels T7 and T8) and eye-blinks or movements (in channels Fp1 and Fp2) were automatically detected using a combination of z-value cutoffs and filtering. Trials in which artifacts were detected were removed.

For each trial, data were bandpass filtered between 1 and 70 Hz and the power spectral density was estimated using Welch’s method with a window length of 4 s and 50% overlap, giving a frequency resolution of 0.25 Hz. We computed relative power, by expressing the power at each frequency resolution point as a fraction of the total power between 1 and 70 Hz. Data were then bandpassed into the delta (1–4 Hz) and alpha (8–13 Hz) frequency bands. The peak power (maximum power) in each trial was computed for each subject and averaged across trials per block.

### pCASL

Perfusion data was acquired using a pCASL sequence from JJ Wang (University of Southern California) with a segmented (2-shot) 3DGRASE readout (Kilroy et al., 2014): labeling duration (LD) = 1800 ms, post-labeling delay (PLD) = 2100 ms, repetition time (TR)/echo time (TE) = 4500/17.32 ms, Turbo/EPI factor = 27/31, field of view = 256 × 256 mm^2^, acquired voxel size = 4.0 × 4.0 × 5.5 mm (24 slices), reconstructed at 2.0 × 2.0 × 5.5 mm. The post-labelling delay was chosen, following plausibility checks, to account for the expected increase in bolus arrival time associated with hyperventilation. To further correct pCASL data for distortion, an additional M0 scan with reversed phase encoding was collected.

For the first hyperventilation task and the first 18 participants, the pCASL exam included 13, 10, 7, 10, 7, 10, 9 tag/control pairs for pre-HV, HV1, rest1, HV2, rest2, HV3, post-HV, respectively. For the second hyperventilation task and the last 13 participants, the pCASL exam included 13, 13, 10, 14, 10, 5, 9 tag/control pairs for pre-HV, HV1, rest1, HV2, rest2, HV3, post-HV respectively. For both protocols, the post-HV block began 145±17 seconds after the last hyperventilation task.

Controls and tags images were separated and rigidly corrected for motion using the first control or tag image (Jenkinson, 2002). After realignment, tags and controls dynamics were re-merged for further pCASL quantification.

To correct for arterial artifact caused by a delay in the transit of blood to the capillaries and leading to hyperintense signal within the arteries, we used an in-house program implemented in Matlab (R2020b, The MathWorks Inc., Natick, MA). The tag-control difference dynamics were computed and any voxel with a z-score greater or lower than 3.5 was excluded from further analysis.

Quantitative perfusion maps were computed using Oxford ASL tools (v4.0.16, http://fsl.fmrib.ox.ac.uk/fsl/fslwiki/oxford_asl) (Chappell et al., 2009) with distortion correction implementation (Andersson et al., 2003).

Using the CSF probability map as a reference, M0 images were used to realign and resample pCASL data to each participant’s anatomical 3D-MRI space. Perfusion maps were sampled using the cortical gray matter, basal ganglia, and thalamus labels.

### Statistical analyses

To show that the hyperventilation protocols were equivalent in their effect on whole-brain CBF, we first compared whole cortical gray matter CBF between the two protocols using non-parametric Mann-Whitney U tests for each HV and rest block.

We then compared baseline values (i.e. pre-HV) for respiratory rates, CBF, and EEG alpha and delta frequency band power between groups using non-parametric Mann-Whitney U tests.

To assess group differences due to HV, we normalised all measurements (i.e. respiratory rate, CBF and EEG band power) by pre-HV values for each individual (i.e. divided by pre-HV values, multiplied by 100%) and pooled across all HV and rest blocks. Each measure was then compared between patient and control groups with the non-parametric Mann-Whitney U test.

The same measures were compared between participants who had epileptiform discharges (ED) evident on their intra-scan EEG recordings (epilepsy-EDs) and those who did not (epilepsy-no-EDs and controls) with the non-parametric Mann-Whitney U test.

Pooling across all rest and HV blocks, correlations between measures were assessed with Spearman’s rank-order correlation tests. Spearman’s rho values are expressed with 0.95 Confidence Intervals [lower limit; upper limit].

All statistical analysis were performed using The jamovi project (2022). *jamovi* (Version 2.2).

## Results

### Participants

Table 1 summarizes participant demographic data. There was no statistical difference between groups for gender (p = 0.879), age (p = 0.631) and BMI (p=0.115). BDI-II (Beck Depression Inventory) score was significantly higher in patients (p=0.043). Individual patient characteristics are given in Table 2. Age at onset was 11.0 ± 4.0 years (range, 5-39); duration of epilepsy 19 ± 18 (0 – 46) years; the interictal interval was 7 ± 53 days (0 – 365). Five of the 13 patients had epileptiform discharges during the scan; two indicated having had absences. Three patients were not taking antiseizure medication, three were on monotherapy, and seven were on a combination of antiseizure drugs.

**Table 1.**
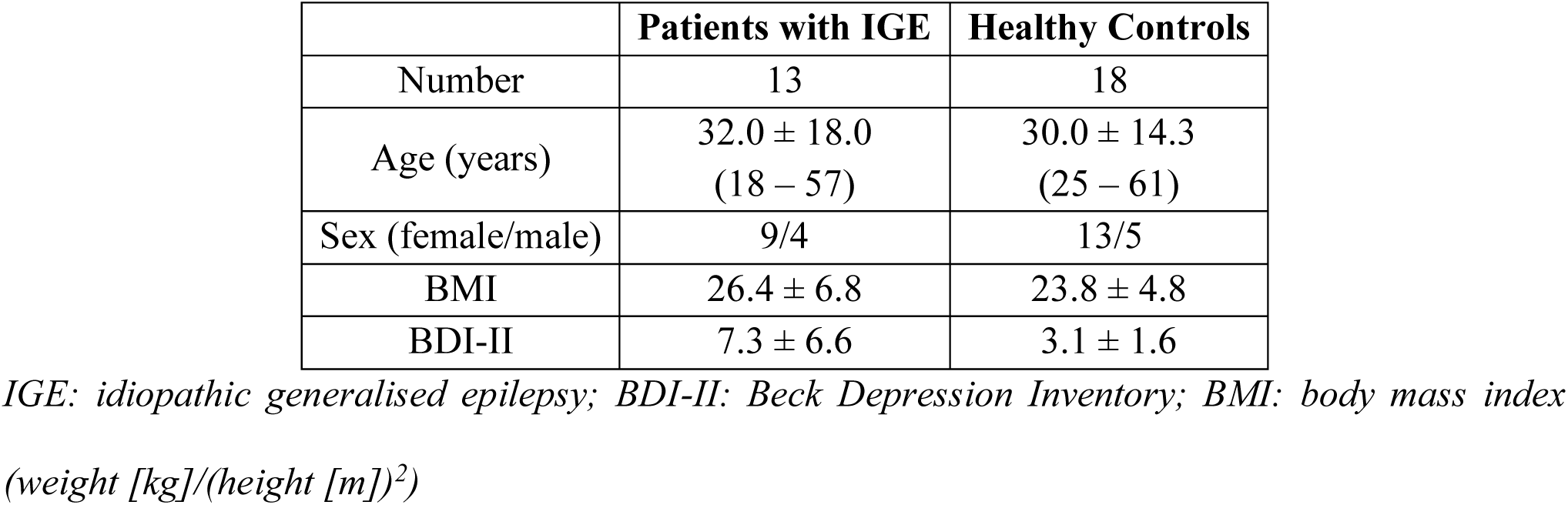
Participant demographic and patient clinical data. Age and age at epilepsy onset are given as median ± interquartile range.

**Table 2.**
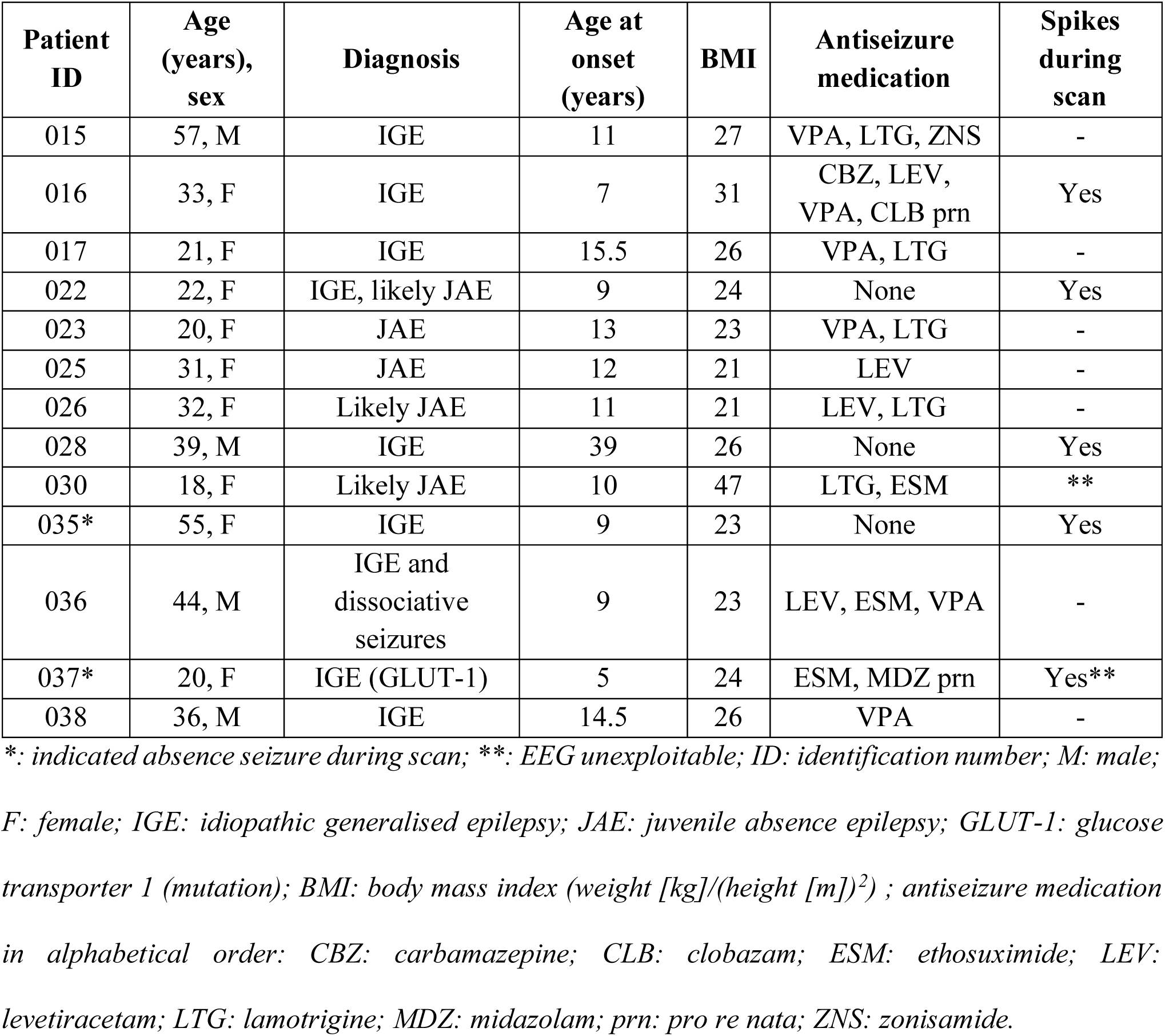
Individual patient data.

### Missing data

Due to technical issues in the data acquisition and/or processing, two pCASL datasets (one control and one IGE patient), two EEG recordings (two patients) and two respiratory rates (two controls) were unexploitable. In the following analyses, pCASL results are reported for n=29 participants (n=17 controls and n=12 IGE patients); EEG results for n=29 participants (n=18 controls and n=11 IGE patients) and respiratory rates for n=29 participants (n=16 controls and n=13 IGE patients).

### Comparison of first and second hyperventilation protocols & baseline data

In the whole cohort of participants (n = 29), by sampling the whole cortical gray matter CBF, no significant difference between the first and the second hyperventilation task protocols was found (p = 0.540 for pre-HV, p = 0.744 for HV1, p = 0.283 for rest1, p = 0.303 for HV2, p = 0.679 for rest2, p = 0.647 for HV3 and p = 0.283 for post-HV).

Baseline pre-HV respiratory rates were not significantly different between patients (median ± interquartile range = 14.5 ± 3.0 respirations/min) and controls (14.13 ± 4.6, p = 0.775).

Baseline pre-HV EEG parameters were also similar between groups, with no significant differences between patients and controls in 1) alpha peak power (patients (median ± interquartile range) = 0.028 ± 0.17 a.u.; controls 0.028 ± 0.16 a.u., p = 0.740), 2) delta peak power (patients (median ± interquartile range) = 0.079 ± 0.22 a.u.; controls 0.057 ± 0.26 a.u., p = 0.09) and 3) alpha/delta ratio peak power (patients (median ± interquartile range) = 0.44 ± 0.25 a.u.; controls 0.47 ± 0.35 a.u., p = 0.276).

Finally, baseline pre-HV CBF parameters also showed no significant differences between patients and controls for 1) cortical gray matter CBF (patients (median ± interquartile range) = 52.6 ± 15.9 mL/min/100g; controls 46.3 ± 8.7 mL/100g/min, p = 0.073), 2) thalamic CBF (patients (median ± interquartile range) = 42.4 ± 12.6 mL/min/100g; controls 41.1 ± 6.6 mL/100g/min, p = 0.245), and 3) basal ganglia CBF (patients (median ± interquartile range) = 33.9 ± 13.6 mL/min/100g; controls 29.4 ± 5.9 mL/100g/min, p = 0.195).

As there were no significant group differences in respiration rates, EEG parameters, or CBF during the baseline pre-HV block, the results for the remainder of the study are given as normalised by the pre-HV value.

### Normalised respiratory rates

Figure 2A shows the normalised respiratory rates in all rest and HV blocks for each group. Normalised respiratory rates increased during HV in both groups, as expected. Respiratory rates decreased below baseline during rest blocks in both groups, with patients showing less of a decrease (i.e. higher normalised respiratory rates during rest) than healthy controls (patients (median ± interquartile range) = 89.7 ± 56.4; controls 74.4 ± 32.8; p = 0.032; Fig. 2). There was no difference in respiratory rates between groups during HV, indicating that both groups performed the hyperventilation task equally well.

**Figure 2.**
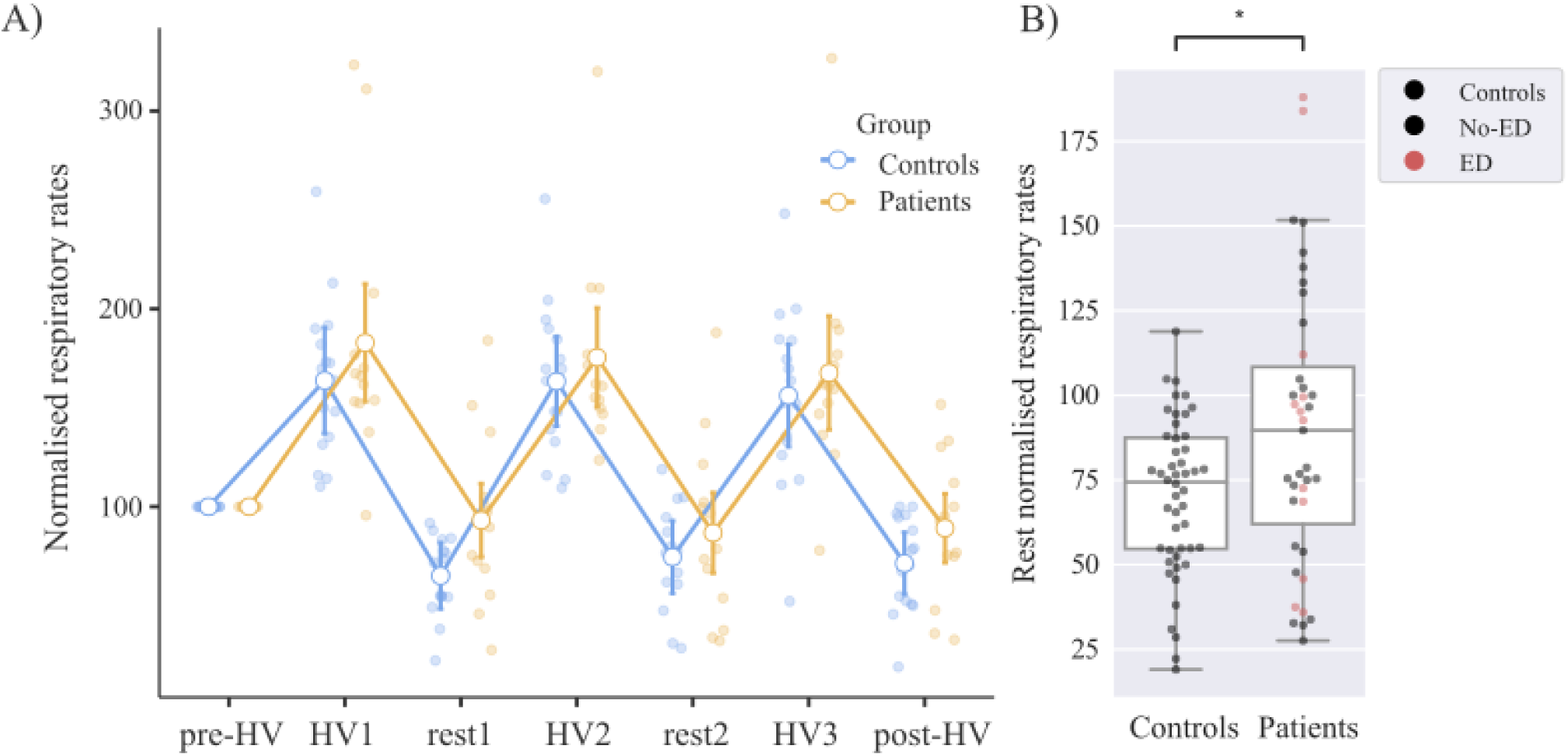
**(A)** Normalised respiratory rates (expressed as a percentage of pre-HV values) in IGE patients (n = 13) and healthy controls (n = 16) for each HV and rest block (white circles represent the mean; error bars are 95% confidence intervals) and **(B)** pooled across rest blocks (square is the mean, horizontal line is the median and error bars are 1.5*IQR). HV: hyperventilation, IGE: idiopathic generalised epilepsy; IQR: interquartile range

### Normalised cortical grey matter and basal ganglia CBF decrease induced by hyperventilation

Figure 3A shows an example of pCASL CBF maps in a control participant. CBF levels clearly decreased throughout the whole brain during the first hyperventilation task (HV1) as compared to pre-HV baseline values, by 35-40% during the hyperventilation tasks (HV1, HV2, and HV3) compared to pre-HV (Figure 3B). CBF remained reduced during the short rest periods between HV tasks and had still not entirely returned to baseline in the post-HV period (145 seconds after the end of HV3). Comparing the two groups, during HV, there was no difference in the decrease in CBF in the cortical grey matter (−37.1 ± 1.3% in controls and −37.9 ± 2.0% in patients, p=0.521) and no difference in the decrease in CBF in the basal ganglia (−35.6 ± 2.0% in controls and −33.3 ± 2.3% in patients, p=0.701).

**Figure 3.**
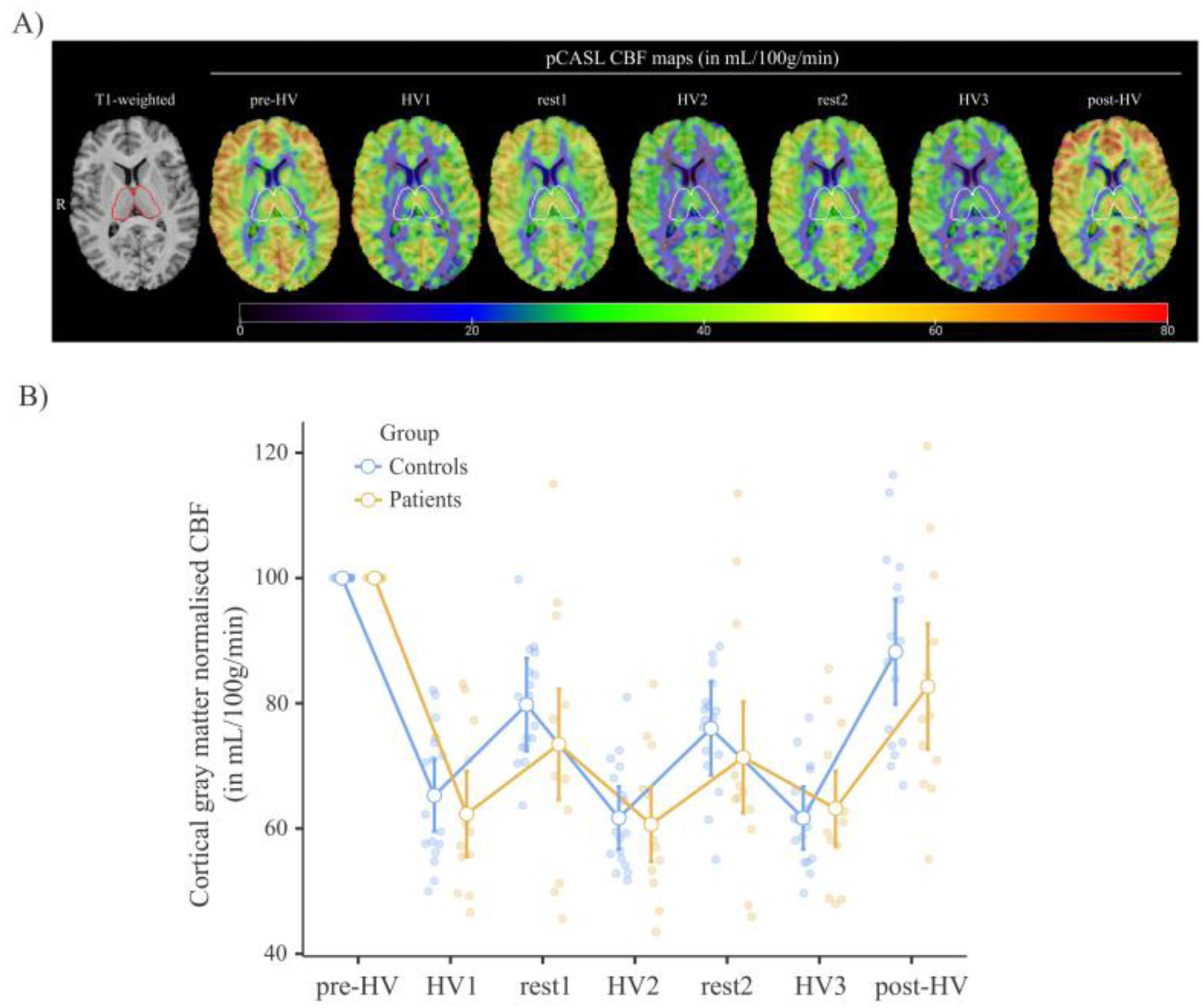
**(A)** illustration of whole-brain CBF maps derived from the pCASL analysis in one control participant. Quantitative CBF maps (in mL/100g/min) are displayed together with the T1 weighted MRI scan. The thalamic ROI is overlain **(B)** Plots of individual normalised CBF values (expressed as percentage of pre-HV value) in the cortical gray matter label in IGE patients (n = 12) and controls (n = 17) at all blocks of the hyperventilation task protocol. Dots represent the mean and bars 95% of the confidence interval. CBF: cerebral blood flow; HV: hyperventilation, IGE: idiopathic generalised epilepsy; MRI: magnetic resonance imaging; pCASL: pseudo-continuous Arterial Spin Labelling

### Normalised thalamic CBF decrease induced by hyperventilation

Figure 4A shows the normalised thalamic CBF in all rest and HV blocks for each group. Like whole-brain patterns of CBF, normalised thalamic CBF decreased relative to baseline throughout the HV tasks and rest periods in both groups, with a greater decrease relative to baseline seen during HV than rest blocks. During the HV blocks, normalised thalamic CBF differed between patients with IGE and healthy controls (p = 0.011), with a decrease relative to baseline of about 35% in healthy controls and 27% in patients with IGE (Fig. 4B). In contrast, there was no group difference in normalised thalamic CBF during the rest blocks.

**Figure 4.**
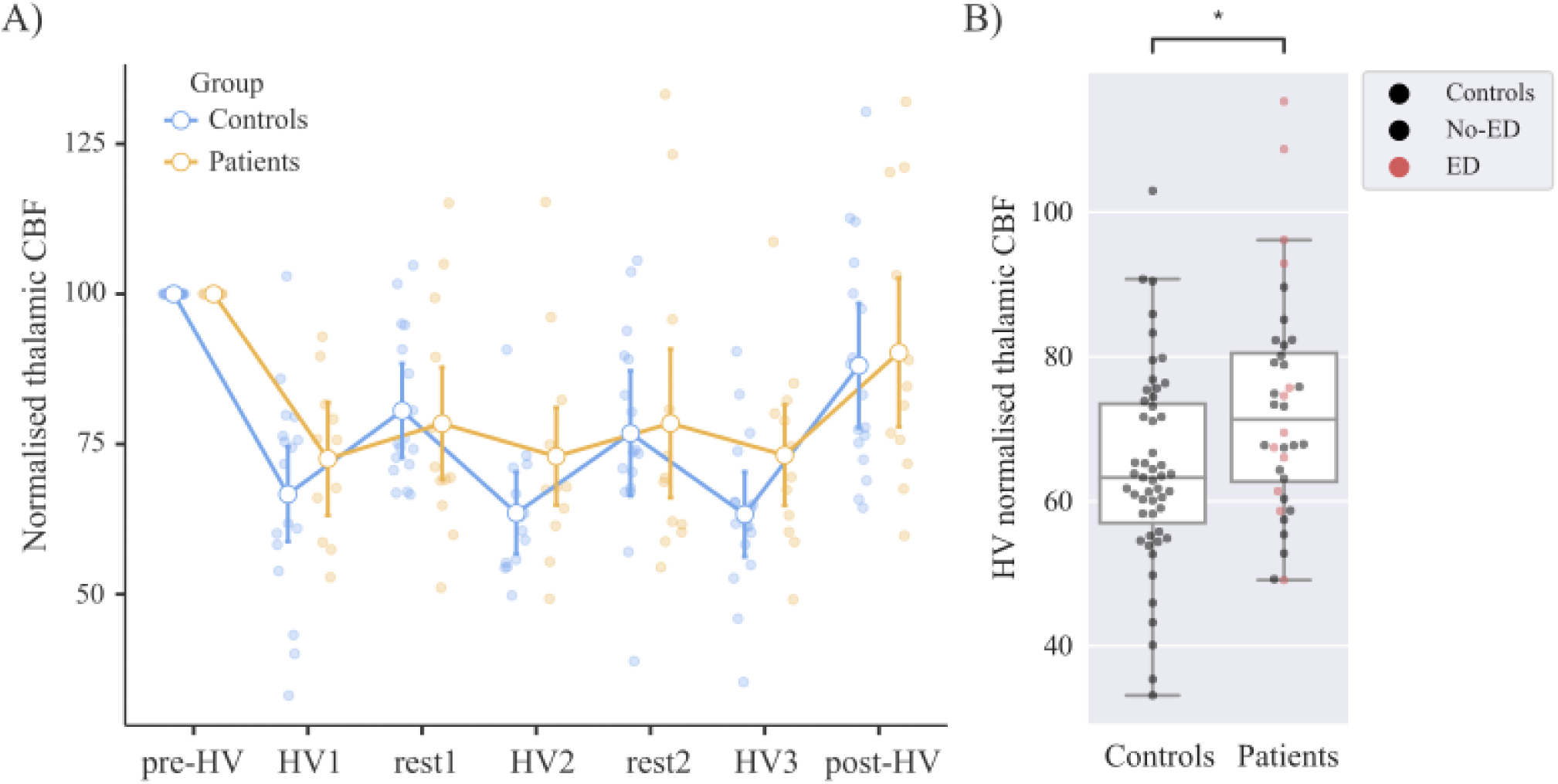
**(A)** Normalised thalamic CBF (expressed as a percentage of pre-HV values) in IGE patients (n = 12) and healthy controls (n = 17) for each HV and rest block (circles represent the mean and error bars are 95% confidence interval) and **(B)** pooled across rest blocks (square is the mean, horizontal line is the median and error bars are 1.5*IQR). *: significant at p < 0.05, CBF: cerebral blood flow; HV: hyperventilation, IGE: idiopathic generalised epilepsy; IQR: interquartile range; MRI: magnetic resonance imaging; pCASL: pseudo-continuous Arterial Spin Labelling

### Normalised EEG band power

There were no significant group differences in the normalised EEG alpha band power either during rest or HV. EEG alpha band power decreased relative to baseline during HV blocks and returned to approximately baseline levels during rest periods in both groups (Fig. 5).

**Figure 5.**
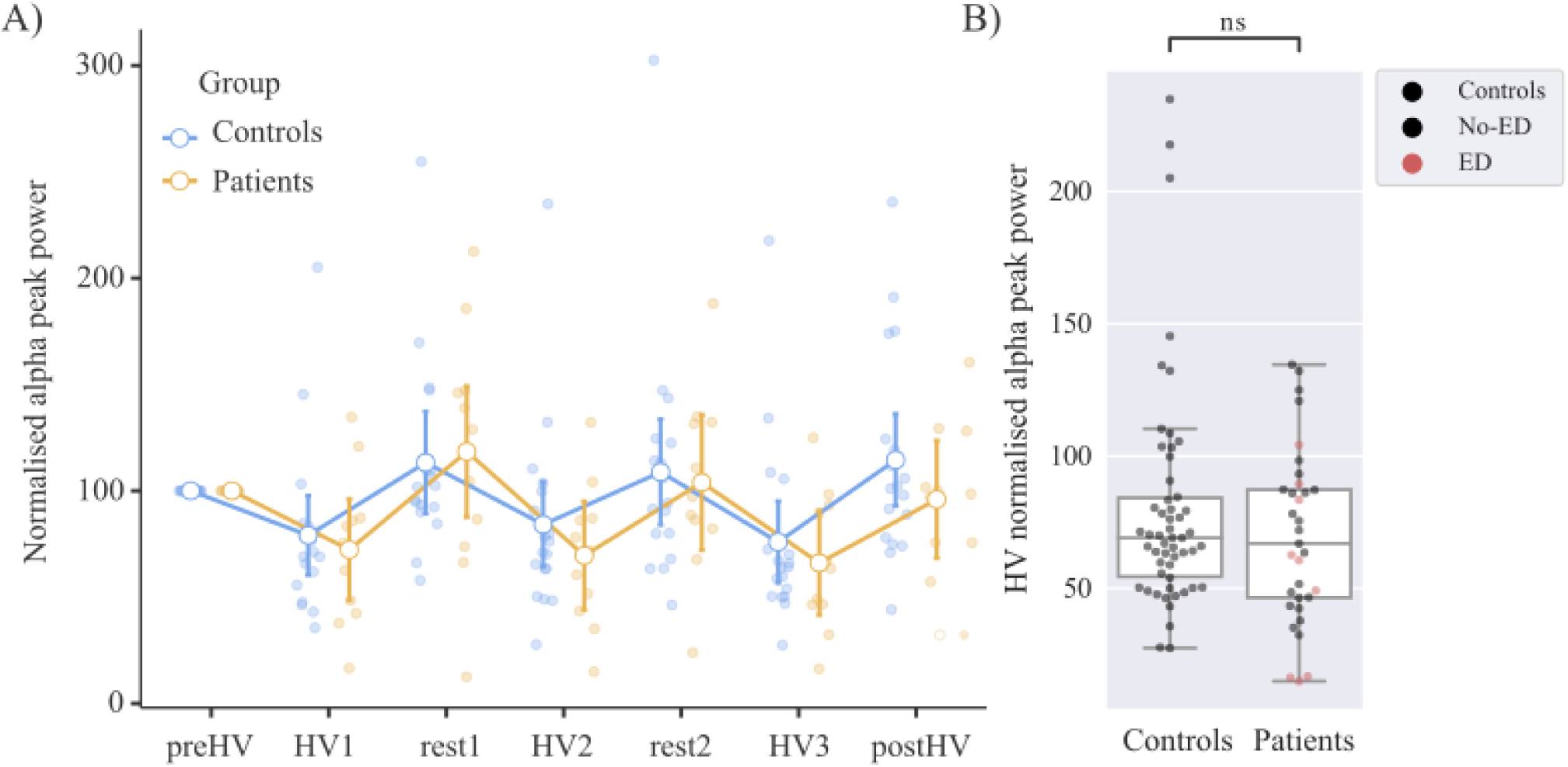
Normalised EEG alpha frequency band power (expressed as a percentage of pre-HV values) in IGE patients (n = 11) and healthy controls (n = 18) for each HV and rest block (circles represent the mean and error bars are 95% confidence interval; difference not significant). HV: hyperventilation, IGE: idiopathic generalised epilepsy; EEG: electroencephalography

Figure 6A shows the normalised EEG delta power averaged across all electrodes in all rest and HV blocks for each group. As expected, EEG delta band power increased during HV blocks relative to baseline and returned approximately to baseline levels during rest blocks in both groups. During HV, EEG delta band power differed between groups (p = 0.045), where patients with IGE showed a smaller increase in delta band power relative to baseline (27%) than healthy controls (42%; Fig. 6B). Figure 6C shows the topographical spread of median normalised EEG delta band power during HV seen over predominantly centro-parietal electrodes. There was no group difference in EEG delta band power during rest.

**Figure 6.**
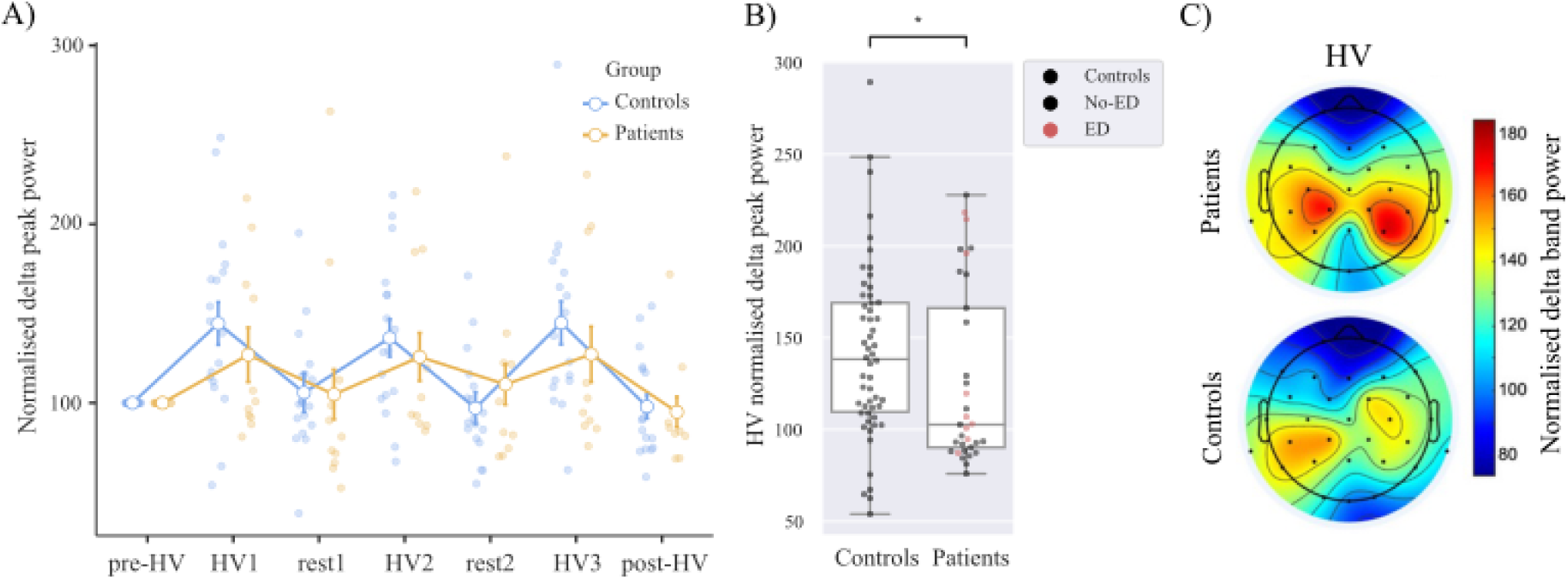
**(A)** Normalised EEG delta frequency band power (expressed as a percentage of pre-HV values) in IGE patients (n = 11) and healthy controls (n = 18) for each HV and rest block (circles represent the mean and error bars are 95% confidence interval) and **(B)** pooled across HV blocks (square is the mean, horizontal line is the median and error bars are 1.5*IQR). **(C)** Topographical plots of the median normalised EEG delta frequency band power during HV blocks. *: significant at p < 0.05, CBF: cerebral blood flow; HV: hyperventilation, IGE: idiopathic generalised epilepsy; IQR: interquartile range; EEG: electroencephalography

### Correlations between measures

Pooling all rest blocks, normalised thalamic CBF was negatively correlated with normalised respiratory rate in both healthy controls (rho = −0.51 [-0.698; −0.255], p < 0.001) and patients with IGE (rho = −0.38 [-0.629; −0.059], p = 0.002) (Fig. 7).

**Figure 7.**
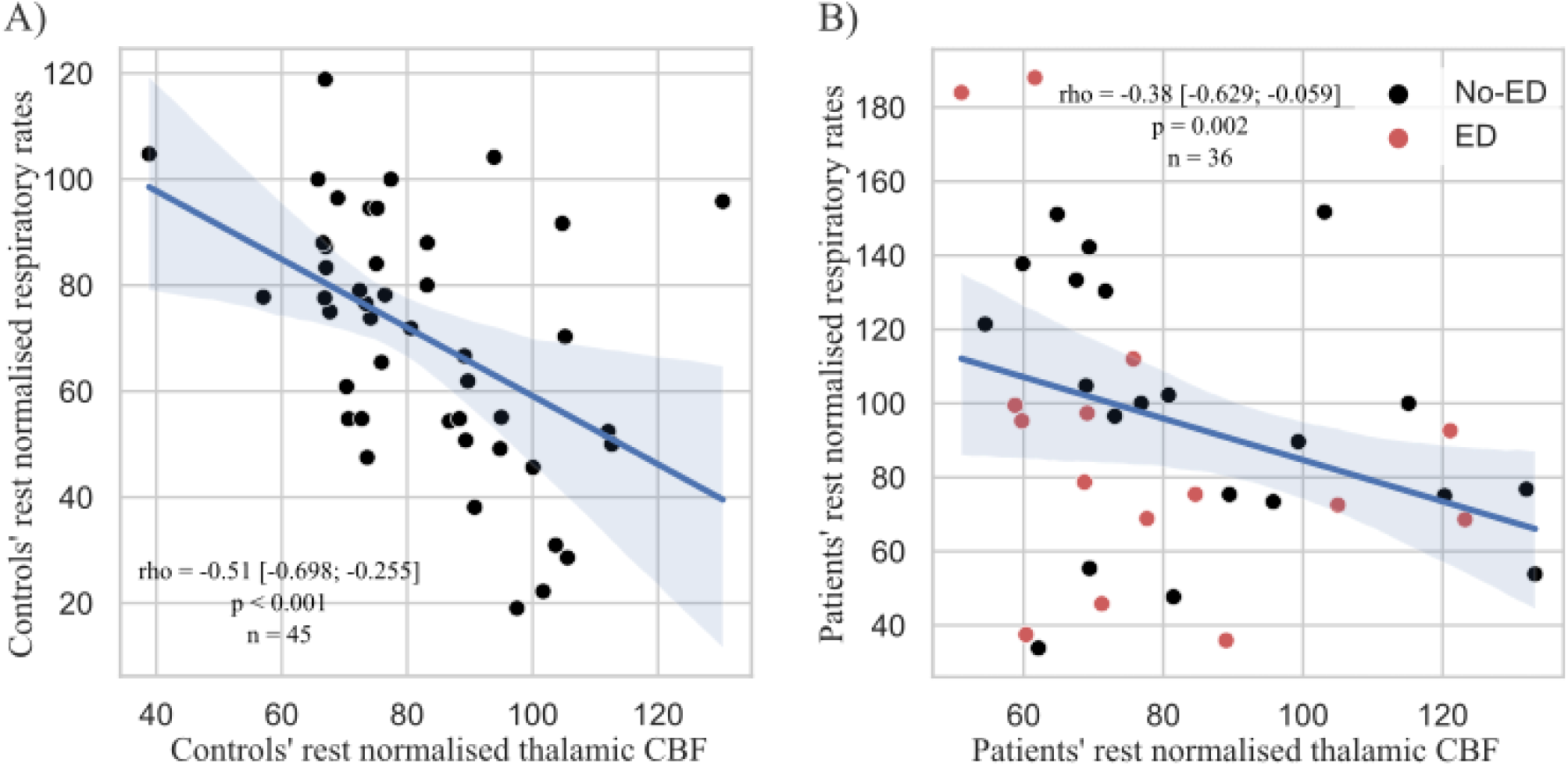
Correlations between normalised thalamic CBF and normalised respiratory rates at rest in healthy controls (A, n = 45 data points from 15 healthy controls) and IGE patients (B, n = 36 data points from 12 IGE patients). Rho values are expressed with 0.95 Confidence Intervals [lower limit; upper limit]. CBF: cerebral blood flow; IGE - idiopathic generalised epilepsy

When pooling all rest and HV blocks, delta frequency band power correlated with thalamic CBF in healthy controls (rho = −0.21 [-0.388; −0.017], p = 0.037), but not in patients with IGE (rho = −0.1 [- 0.345; 0.157], p = 0.59) (Fig. 8). EEG band power and respiratory rates were not significantly correlated.

**Figure 8.**
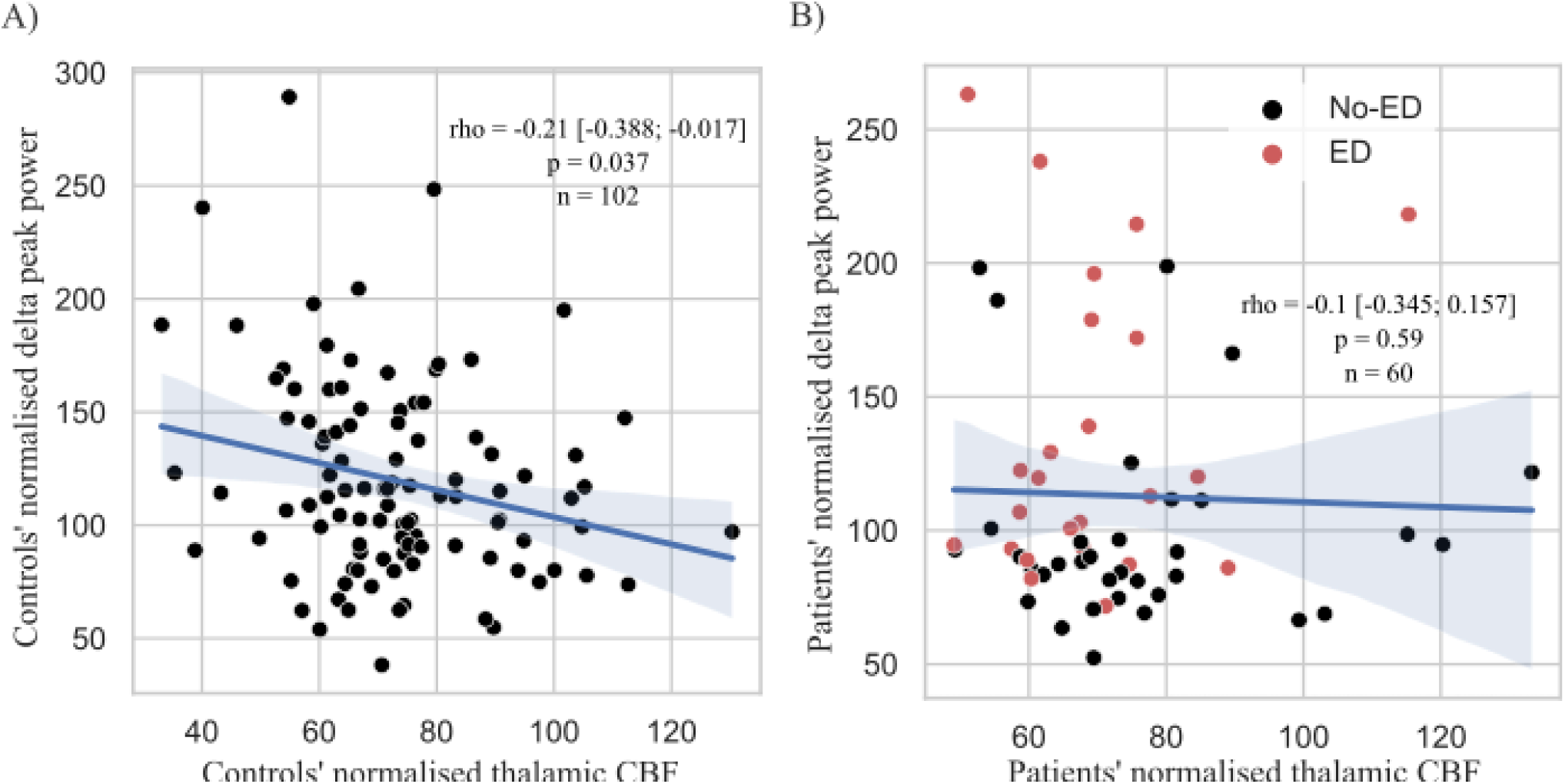
Correlation between normalised thalamic CBF and normalised EEG delta frequency band peak power in both rest and HV blocks in healthy controls (A, n = 102 data points from 17 healthy controls) and IGE patients (B, n = 102 data points from 17 healthy controls). Rho values are expressed with 0.95 Confidence Intervals [lower limit; upper limit]. CBF: cerebral blood flow; EEG: electroencephalography; IGE: idiopathic generalised epilepsy

### Comparisons between spiking and non-spiking patients

Comparing the participants with exploitable EEG results (four spiking patients, seven non-spiking patients), spiking patients had 1) lower mean alpha peak power during HV, 2) higher delta peak power during rest, 3) a lower alpha/delta peak power during rest (differences not significant after correction for multiple comparisons; data not shown).

## Discussion

To our knowledge, this is the first study to simultaneously investigate changes in CBF and EEG band power due to HV in patients with IGE compared with healthy controls.

Our main novel findings were that during HV, patients showed a significantly smaller decrease of normalised thalamic CBF (25% vs 35% in controls) and also had a significantly smaller increase in EEG delta frequency band power (27.2% vs 41.9%). There was also a weaker negative correlation between respiratory rate and thalamic CBF in patients in contrast to controls, and in contrast to controls there was no negative correlation between thalamic CBF and delta band power in patients.

Respiratory rates were not significantly different between groups prior to all tasks and during HV, indicating that both groups performed the task well. Similarly, global, basal ganglia, and thalamic CBF decreased substantially in both patients and controls, in line with previous studies showing that hyperventilation significantly reduces CBF (Blinn & Noell, 1949; Ito et al., 2002; Kety & Schmidt, 1946). Together, these findings indicate that the hyperventilation task worked as intended. Patients had overall slightly higher respiratory rates which might be linked to lower cardiorespiratory fitness which is frequently found across the epilepsies (Steinhoff et al., 1996; Vancini et al., 2015).

Multiple lines of enquiry link the thalamus to IGE. The thalamus has been found hypoperfused in patients with IGE interictally (Sone et al., 2017), and thalamocortical circuits are implicated in absence seizure generation and spread (Aghakhani et al., 2004; Avoli, 2012). Importantly, we found no significant differences between patients and controls in CBF, or in CBF changes during hyperventilation, when examining either all of the gray matter, or structures of a similar size, i.e. the basal ganglia. This suggests that thalamic blood flow reactivity to HV is selectively altered in IGE.

There is a known correlation between decreased regional blood flow (including in the thalamus) and increased EEG delta power in healthy individuals during slow wave sleep (Davis et al., 2011; Hofle et al., 1997). The lack of correlation between thalamic CBF and delta band power in patients in this study could therefore suggest that patients do not generate enough delta power to decrease thalamic cerebral blood flow, or they lack the mechanism linking delta generation to decreased blood flow. An alternative explanation is that the primary defect is with the cerebral autoregulation of blood flow in the thalamus, i.e. patients do not decrease their thalamic blood flow enough to then allow the generation of delta waves visible on scalp EEG.

Our study has several limitations. In the majority of patients, HV did not activate spike-wave discharges on their EEG and only two indicated brief absences. In contrast, in the study of hyperventilation-induced endogenous opioid release by Bartenstein et al. (Bartenstein et al., 1993), clinical absences were triggered for substantial fractions of the hyperventilation task. This may be due to optimized treatment since the 1993 study: of their eight patients, four (50%) were treated with sodium channel blockers (phenytoin, carbamazepine) now widely accepted to carry a risk of seizure aggravation in IGE (Chaves & Sander, 2005; Somerville, 2009; Thomas et al., 2006). Similarly, in Prevett et al. (Prevett et al., 1995), five of eight patients were treated with sodium channel blockers and in all of those spike-wave activity was triggered by hyperventilation (5-42% of the time of the ictal scan), whereas only one of their three patients on valproate monotherapy had ictal discharges. Interestingly, the only patient in our cohort whose antiseizure medication included carbamazepine was one of five with epileptiform discharges during the scan. In addition, even though we only recruited patients with very frequent and/or hyperventilation-inducible absence seizures, only five of the 13 patients had EEG discharges and only two indicated absence seizures. Similar effects of “EEG normalisation” in a scanner environment have been seen for patients with focal epilepsies (e.g. (Salek-Haddadi et al., 2006)).

Possibly due to the better contemporary treatment of adults with HV-inducible absences compared to the earlier studies (Bartenstein et al., 1993; Prevett et al., 1995), recruitment proved harder than expected and was curtailed by the restrictions imposed by the COVID-19 pandemic, meaning we did not achieve recruitment of all 18 patients planned. While there is a risk of type II error, we were able to show significant and coherent differences between the patients and controls.

Different thalamic nuclei may have different roles in absence initiation and maintenance (Lüttjohann & van Luijtelaar, 2022). Due to the low signal-to-noise ratio and relatively low spatial resolution inherent in the pCASL measurements, we did not attempt to further parcellate thalamic subnuclei.

It was also not possible due to experimental constraints to measure pCO_2_ directly; however, respiratory rates during HV were very well aligned with the acoustic prompt and not different between groups.

## Conclusions

This first simultaneous study of hyperventilation-induced EEG band power and regional CBF changes has demonstrated regionally specific reductions in cerebral autoregulation in the thalamus of patients with IGE, accompanied by and correlated with decreases in scalp EEG delta band activation. Further studies are necessary to explore the mechanisms underlying the phenomenon, e.g. by pharmacological rather than pCO_2_-mediated manipulation of CBF or simultaneous studies in preclinical models of IGE.

## Data availability statement

Data that support the findings of this study are available upon reasonable request from the corresponding author.

## Abbreviations

ASL: arterial spin labelling
CBF: cerebral blood flow
ECG: electrocardiogram
ED: epileptiform discharges
EEG: electroencephalography
FWHM: full width at half maximum
HV: hyperventilation
IGE: idiopathic generalised epilepsy
IQR: interquartile range
MAPER: multi-atlas propagation with enhanced registration
MNI: Montreal Neurological Institute
MR: magnetic resonance
MRI: magnetic resonance imaging
pCASL: pseudo-continuous arterial spin labelling
pCO_2_: partial pressure of carbon dioxide
pH: potential of hydrogen
RR: respiratory rate

## References

Aghakhani, Y., Bagshaw, a P., Bénar, C. G., Hawco, C., Andermann, F., Dubeau, F., & Gotman, J. (2004). fMRI activation during spike and wave discharges in idiopathic generalized epilepsy. Brain, 127(Pt 5), 1127-1144. 10.1093/brain/awh136

Andersson, J. L. R., Skare, S., & Ashburner, J. (2003). How to correct susceptibility distortions in spin-echo echo-planar images : Application to diffusion tensor imaging. NeuroImage, 20(2), 870-888. 10.1016/S1053-8119(03)00336-7

Avoli, M. (2012). A brief history on the oscillating roles of thalamus and cortex in absence seizures. Epilepsia, 53(5), 779-789. 10.1111/j.1528-1167.2012.03421.x

Bartenstein, P. A., Duncan, J. S., Prevett, M. C., Cunningham, V. J., Fish, D. R., Jones, A. K. P., Luthra, S. K., Sawle, G. V., & Brooks, D. J. (1993). Investigation of the opioid system in absence seizures with positron emission tomography. Journal of Neurology, Neurosurgery & Psychiatry, 56(12), 1295-1302. 10.1136/jnnp.56.12.1295

Berger, H. (1934). Über das Elektrenkephalogramm des Menschen. DMW - Deutsche Medizinische Wochenschrift, 60(51), 1947-1949. 10.1055/s-0028-1130334

Berman, R., Negishi, M., Vestal, M., Spann, M., Chung, M. H., Bai, X., Purcaro, M., Motelow, J. E., Danielson, N., Dix-Cooper, L., Enev, M., Novotny, E. J., Constable, R. T., & Blumenfeld, H. (2010). Simultaneous EEG, fMRI, and behavior in typical childhood absence seizures. Epilepsia, 51(10), 2011-2022. 10.1111/j.1528-1167.2010.02652.x

Blinn, K. A., & Noell, W. K. (1949). Continuous measurement of alveolar CO2 tension during the hyperventilation test in routine electroencephalography. Electroencephalography and Clinical Neurophysiology, 1(1-4), 333-342. 10.1016/0013-4694(49)90198-4

Chappell, M. A., Groves, A. R., Whitcher, B., & Woolrich, M. W. (2009). Variational Bayesian Inference for a Nonlinear Forward Model. IEEE Transactions on Signal Processing, 57(1), 223-236. 10.1109/TSP.2008.2005752

Chaves, J., & Sander, J. W. (2005). Seizure Aggravation in Idiopathic Generalized Epilepsies. Epilepsia, 46(s9), 133-139. 10.1111/j.1528-1167.2005.00325.x

Dai, W., Garcia, D., De Bazelaire, C., & Alsop, D. C. (2008). Continuous flow-driven inversion for arterial spin labeling using pulsed radio frequency and gradient fields : Pulsed Continuous Arterial Spin Labeling. Magnetic Resonance in Medicine, 60(6), 1488-1497. 10.1002/mrm.21790

Davis, C. J., Clinton, J. M., Jewett, K. A., Zielinski, M. R., & Krueger, J. M. (2011). Delta Wave Power : An Independent Sleep Phenotype or Epiphenomenon? Journal of Clinical Sleep Medicine, 7(5 Suppl). 10.5664/JCSM.1346

Delso, G., Fürst, S., Jakoby, B., Ladebeck, R., Ganter, C., Nekolla, S. G., Schwaiger, M., & Ziegler, S. I. (2011). Performance Measurements of the Siemens mMR Integrated Whole-Body PET/MR Scanner. Journal of Nuclear Medicine, 52(12), 1914-1922. 10.2967/jnumed.111.092726

Depaulis, A., & Charpier, S. (2018). Pathophysiology of absence epilepsy : Insights from genetic models. Neuroscience Letters, 667, 53-65. 10.1016/j.neulet.2017.02.035

Elliott, K. A. C., & Jasper, H. H. (1949). Physiological Salt Solutions for Brain Surgery. Journal of Neurosurgery, 6(2), 140-152. 10.3171/jns.1949.6.2.0140

Faillenot, I., Heckemann, R. A., Frot, M., & Hammers, A. (2017). Macroanatomy and 3D probabilistic atlas of the human insula. NeuroImage, 150(December 2016), 88-98. 10.1016/j.neuroimage.2017.01.073

Foerster, O. (1924). Hyperventilationsepilepsie. Z Neurol Psychiatrie, 38, 289-293.

Fojtiková, D., Brázdil, M., Horký, J., Mikl, M., Kuba, R., Krupa, P., & Rektor, I. (2006). Magnetic resonance spectroscopy of the thalamus in patients with typical absence epilepsy. Seizure, 15(7), 533-540. 10.1016/j.seizure.2006.06.007

Gibbs, F. A., Davis, H., & Lennox, W. G. (1968). The Electro Encephalogram in Epilepsy and in Conditions of Impaired Consciousness. American Journal of EEG Technology, 8(2), 59-73. 10.1080/00029238.1968.11080707

Gotman, J., Grova, C., Bagshaw, A., Kobayashi, E., Aghakhani, Y., & Dubeau, F. (2005). Generalized epileptic discharges show thalamocortical activation and suspension of the default state of the brain. Proceedings of the National Academy of Sciences of the United States of America, 102(42), 15236-15240. 10.1073/pnas.0504935102

Gousias, I. S., Rueckert, D., Heckemann, R. A., Dyet, L. E., Boardman, J. P., Edwards, A. D., & Hammers, A. (2008). Automatic segmentation of brain MRIs of 2-year-olds into 83 regions of interest. NeuroImage, 40(2), 672-684. 10.1016/j.neuroimage.2007.11.034

Hammers, A., Allom, R., Koepp, M. J., Free, S. L., Myers, R., Lemieux, L., Mitchell, T. N., Brooks, D. J., & Duncan, J. S. (2003). Three-dimensional maximum probability atlas of the human brain, with particular reference to the temporal lobe. Human Brain Mapping, 19(4), 224-247. 10.1002/hbm.10123

Heckemann, R. A., Keihaninejad, S., Aljabar, P., Rueckert, D., Hajnal, J. V., & Hammers, A. (2010). Improving intersubject image registration using tissue-class information benefits robustness and accuracy of multi-atlas based anatomical segmentation. NeuroImage, 51(1), 221-227. 10.1016/j.neuroimage.2010.01.072

Heckemann, R. A., Ledig, C., Gray, K. R., Aljabar, P., Rueckert, D., Hajnal, J. V., & Hammers, A. (2015). Brain Extraction Using Label Propagation and Group Agreement : Pincram. PLOS ONE, 10(7), e0129211. 10.1371/journal.pone.0129211

Hirsch, E., French, J., Scheffer, I. E., Bogacz, A., Alsaadi, T., Sperling, M. R., Abdulla, F., Zuberi, S. M., Trinka, E., Specchio, N., Somerville, E., Samia, P., Riney, K., Nabbout, R., Jain, S., Wilmshurst, J. M., Auvin, S., Wiebe, S., Perucca, E.,… Wirrell, E. C. (2022). ILAE definition of the Idiopathic Generalized Epilepsy Syndromes : Position statement by the ILAE Task Force on Nosology and Definitions. Epilepsia, 63(6), 1475-1499. 10.1111/epi.17236

Hirsch, E., French, J., Scheffer, I. E., Bogacz, A., Alsaadi, T., Sperling, M. R., Abdulla, F., Zuberi, S. M., Trinka, E., Specchio, N., Somerville, E., Samia, P., Riney, K., Nabbout, R., Jain, S., Wilmshurst, J. M., Auvin, S., Wiebe, S., Perucca, E.,… Zhou., D. (2022). ILAE definition of the Idiopathic Generalized Epilepsy Syndromes : Position statement by the ILAE Task Force on Nosology and Definitions. Epilepsia, 63(6), 1475-1499. 10.1111/epi.17236

Hofle, N., Paus, T., Reutens, D., Fiset, P., Gotman, J., Evans, A. C., & Jones, B. E. (1997). Regional cerebral blood flow changes as a function of delta and spindle activity during slow wave sleep in humans. Journal of Neuroscience, 17(12), 4800-4808. 10.1523/jneurosci.17-12-04800.1997

Ito, H., Kanno, I., Ibaraki, M., & Hatazawa, J. (2002). Effect of Aging on Cerebral Vascular Response to Paco 2 Changes in Humans as Measured by Positron Emission Tomography. Journal of Cerebral Blood Flow & Metabolism, 22(8), 997-1003. 10.1097/00004647-200208000-00011

Jenkinson, M. (2002). Improved Optimization for the Robust and Accurate Linear Registration and Motion Correction of Brain Images. NeuroImage, 17(2), 825-841. 10.1016/S1053-8119(02)91132-8

Kabay, S. C., Gumustas, O. G., Karaman, H. O., Ozden, H., & Erdinc, O. (2010). A proton magnetic resonance spectroscopic study in juvenile absence epilepsy in early stages. European Journal of Paediatric Neurology, 14(3), 224-228. 10.1016/j.ejpn.2009.06.004

Kety, S. S., & Schmidt, C. F. (1946). The effects of active and passive hyperventilation on cerebral blood flow, cerebral oxygen consumption, cardiac output, and blood pressure of normal young men. Journal of Clinical Investigation, 25(1), 107-119. 10.1172/JCI101680

Kilroy, E., Apostolova, L., Liu, C., Yan, L., Ringman, J., & Wang, D. J. J. (2014). Reliability of two-dimensional and three-dimensional pseudo-continuous arterial spin labeling perfusion MRI in elderly populations : Comparison with 15o-water positron emission tomography: Reliability of 2D Versus 3D pCASL. Journal of Magnetic Resonance Imaging, 39(4), 931-939. 10.1002/jmri.24246

Lüttjohann, A., & van Luijtelaar, G. (2022). The role of thalamic nuclei in genetic generalized epilepsies. Epilepsy Research, 182(February 2022), 106918. 10.1016/j.eplepsyres.2022.106918

Moeller, F., Siebner, H. R., Wolff, S., Muhle, H., Boor, R., Granert, O., Jansen, O., Stephani, U., & Siniatchkin, M. (2008). Changes in activity of striato–thalamo–cortical network precede generalized spike wave discharges. NeuroImage, 39(4), 1839-1849. 10.1016/j.neuroimage.2007.10.058

Nims, L. F., Gibbs, E. L., Lennox, W. G., Gibbs, F. A., & Williams, D. (1940). Adjustment of acid-base balance of patients with petit mal epilepsy to overventilation. Archives of Neurology And Psychiatry, 43(2), 262. 10.1001/archneurpsyc.1940.02280020070005

Oostenveld, R., Fries, P., Maris, E., & Schoffelen, J.-M. (2011). FieldTrip : Open Source Software for Advanced Analysis of MEG, EEG, and Invasive Electrophysiological Data. Computational Intelligence and Neuroscience, 2011, 1-9. 10.1155/2011/156869

Pardoe, H., Pell, G. S., Abbott, D. F., Berg, A. T., & Jackson, G. D. (2008). Multi-site voxel-based morphometry : Methods and a feasibility demonstration with childhood absence epilepsy. NeuroImage, 42(2), 611-616. 10.1016/j.neuroimage.2008.05.007

Pavilla, A., Arrigo, A., Mejdoubi, M., Duvauferrier, R., Gambarota, G., & Saint-Jalmes, H. (2018). Measuring Cerebral Hypoperfusion Induced by Hyperventilation Challenge With Intravoxel Incoherent Motion Magnetic Resonance Imaging in Healthy Volunteers. Journal of Computer Assisted Tomography, 42(1), 85-91. 10.1097/RCT.0000000000000640

Prevett, M. C., Duncan, J. S., Jones, T., Fish, D. R., & Brooks, D. J. (1995). Demonstration of thalarnic activation during typical absence seizures using H2 15O and PET. Neurology, 45(7), 1396-1402. 10.1212/WNL.45.7.1396

Rosett, J. (1924). The experimental production of rigidity, of abnormal involuntary movements and of abnormal states of consciousness in man. Brain, 47, 293-336.

Salek-Haddadi, A., Diehl, B., Hamandi, K., Merschhemke, M., Liston, A., Friston, K., Duncan, J. S., Fish, D. R., & Lemieux, L. (2006). Hemodynamic correlates of epileptiform discharges : An EEG-fMRI study of 63 patients with focal epilepsy. Brain Research, 1088(1), 148-166. 10.1016/j.brainres.2006.02.098

Somerville, E. R. (2009). Some treatments cause seizure aggravation in idiopathic epilepsies (especially absence epilepsy). Epilepsia, 50, 31-36. 10.1111/j.1528-1167.2009.02233.x

Sone, D., Watanabe, M., Ota, M., Kimura, Y., Sugiyama, A., Maekawa, T., Okura, M., Enokizono, M., Imabayashi, E., Sato, N., & Matsuda, H. (2017). Thalamic hypoperfusion and disrupted cerebral blood flow networks in idiopathic generalized epilepsy : Arterial spin labeling and graph theoretical analysis. Epilepsy Research, 129, 95-100. 10.1016/j.eplepsyres.2016.12.009

Steinhoff, B. J., Neusiiss, K., Thegeder, H., & Reimers, C. D. (1996). Leisure Time Activity and Physical Fitness in Patients with Epilepsy. Epilepsia, 37(12), 1221-1227. 10.1111/j.1528-1157.1996.tb00557.x

Tangwiriyasakul, C., Perani, S., Centeno, M., Yaakub, S. N., Abela, E., Carmichael, D. W., & Richardson, M. P. (2018). Dynamic brain network states in human generalized spike-wave discharges. Brain, 141(10), 2981-2994. 10.1093/brain/awy223

Thomas, P., Valton, L., & Genton, P. (2006). Absence and myoclonic status epilepticus precipitated by antiepileptic drugs in idiopathic generalized epilepsy. Brain, 129(5), 1281-1292. 10.1093/brain/awl047

Van der Worp, H. B., Kraaier, V., Wieneke, G. H., & Van Huffelen, A. C. (1991). Quantitative EEG during progressive hypocarbia and hypoxia. Hyperventilation-induced EEG changes reconsidered. Electroencephalography and Clinical Neurophysiology, 79(5), 335-341. 10.1016/0013-4694(91)90197-C

Vancini, R. L., Lira, C. A. B. de, Andrade, M. dos S., Lima, C. de, & Arida, R. M. (2015). Low levels of maximal aerobic power impair the profile of mood state in individuals with temporal lobe epilepsy. Arquivos de Neuro-Psiquiatria, 73(1), 7-11. 10.1590/0004-282X20140188

Vaudano, A. E., Laufs, H., Kiebel, S. J., Carmichael, D. W., Hamandi, K., Guye, M., Thornton, R., Rodionov, R., Friston, K. J., Duncan, J. S., & Lemieux, L. (2009). Causal hierarchy within the thalamo-cortical network in spike and wave discharges. PloS one, 4(8), e6475. 10.1371/journal.pone.0006475

Wild, H. M., Heckemann, R. A., Studholme, C., & Hammers, A. (2017). Gyri of the human parietal lobe : Volumes, spatial extents, automatic labelling, and probabilistic atlases. PLOS ONE, 12(8), e0180866. 10.1371/journal.pone.0180866

Wu, W.-C., Fernández-Seara, M., Detre, J. A., Wehrli, F. W., & Wang, J. (2007). A theoretical and experimental investigation of the tagging efficiency of pseudocontinuous arterial spin labeling. Magnetic Resonance in Medicine, 58(5), 1020-1027. 10.1002/mrm.21403

Yaakub, S. N., Tangwiriyasakul, C., Abela, E., Koutroumanidis, M., Elwes, R. D. C., Barker, G. J., & Richardson, M. P. (2020). Heritability of alpha and sensorimotor network changes in temporal lobe epilepsy. Annals of Clinical and Translational Neurology, 7(5), 667-676. 10.1002/acn3.51032

Yamatani, M., Konishi, T., Murakami, M., & Okuda, T. (1995). Hyperventilation activation on EEG recording in children with epilepsy. Pediatric Neurology, 13(1), 42-45. 10.1016/0887-8994(95)00089-X

